# Integrin loss and tissue mechanics trigger progressive disruption of simple epithelial

**DOI:** 10.64898/2026.04.17.719173

**Authors:** Lourdes Rincón-Ortega, Cecilia H. Fernández-Espartero, Isabel M. Palacios, Acaimo González-Reyes, María D. Martín-Bermudo

## Abstract

Simple epithelia form cohesive sheets anchored to a basement membrane (BM), yet the mechanisms that preserve their monolayered architecture remain poorly understood. Here, we address this knowledge gap using the simple follicular epithelium of *Drosophila* as a model. Combining live imaging, quantitative image analysis, manipulation of BM mechanical properties and biophysical measurements, our results provide evidence supporting the role of integrins in orienting cell division *in vivo*. They also reveal two previously unrecognized integrin functions essential for epithelial integrity: promoting timely reintegration of displaced cells following non-planar divisions and modulating junctional tension. These activities underpin a stepwise model of epithelial disruption upon integrin removal. An initial ectopic layer arises from altered division orientation and delayed reintegration. Within this layer, integrin-mutant cells exhibit exacerbated defects in both processes, along with increased junctional tension, which together drive progressive epithelial disruption leading to multilayering. BM mechanical properties further modulate these processes, shaping regional susceptibility to disruption. Together, our work defines how integrin-mediated adhesion and BM mechanics maintain epithelial architecture, while revealing discrete intermediate stages of breakdown with potential relevance to epithelial disorganisation in diseases such as cancer.

## Introduction

Epithelia, one of the four basic tissue types in the body, consist of continuous sheets of cells supported by a basement membrane (BM). They line both internal and external surfaces of all multicellular organisms. Epithelia are broadly classified in two main types: simple, composed of a single layer of cells directly attached to the underlying BM, or stratified, consisting of multiple layers, in which only basal cells contact the BM. Both types are essential and serve distinct yet complementary functions. Simple epithelia primarily mediate absorption and secretion, whereas stratified epithelia generally provide protection against physical and chemical insults and pathogens. The establishment and maintenance of these two architectures are therefore critical for organismal homeostasis. However, while the mechanisms underlying the formation and maintenance of stratified epithelia are relatively well characterized, those governing simple epithelial architecture are less understood.

Epithelial organization is established and maintained through coordinated cellular and mechanical mechanisms. Mitotic spindle orientation is a key regulator of epithelial tissue architecture, as it dictates whether divisions occur parallel to the epithelial plane, preserving a simple monolayered organization (planar divisions), or at an angle, initially displacing daughter cells from the plane, promoting tissue stratification (non-planar divisions). Numerous studies in mammalian—including skin, hair follicle, tongue, and prostate—and non-mammalian epithelia have highlighted the importance of oriented cell divisions for proper epithelial morphogenesis and tissue architecture (Byrd et al., 2016; Fernández-Miñán et al., 2007; Lechler and Fuchs, 2005; Schaeffer et al., 2004; Seldin et al., 2016). However, emerging evidence challenges the view that spindle orientation alone dictates epithelial organization. For example, experiments with skin explant cultures have shown that epidermal stratification can be uncoupled from cell division orientation (Damen et al., 2021). Moreover, in embryonic mouse kidney, zebrafish neuroepithelium and several *Drosophila* tissues, non-planar divisions can be tolerated, as displaced cells reintegrate into the epithelial layer to preserve tissue integrity (Bergstralh et al., 2015; Ciruna et al., 2006; Packard et al., 2013). Together, these findings suggest that mechanisms beyond spindle orientation contribute to establishing and maintaining simple versus stratified tissue architecture.

Cell–BM interactions are critical determinants of epithelial organization. Integrins, heterodimeric transmembrane receptors composed of α and β subunits, are the principal mediators of cell adhesion to the BM and have been implicated in maintaining simple epithelial architecture and suppressing stratification. Downregulation of β1 integrin (ITGB1) is part of the normal stratification program in the human epidermis, and loss of ITGB1 leads to abnormal multilayering in both skin and lung tissues (Brakebusch et al., 2000; Jones and Watt, 1993; Simpson et al., 2011). Furthermore, conditional deletion of β1 integrin in the developing mouse lung epithelium increases the frequency of mitotic spindles oriented perpendicular to the BM, resulting in conversion of a monolayered epithelium into a multilayered structure (Chen and Krasnow, 2012). Despite this evidence, the precise mechanisms by which integrins preserve simple epithelial architecture remain incompletely comprehended.

The follicular epithelium (FE) of the adult *Drosophila* ovary provides a powerful model to investigate these mechanisms. The ovary is composed of 16-18 tubular structures called ovarioles, each containing a germarium at the anterior end and progressively older egg chambers towards the posterior end (Spradling, 1993). Each egg chamber consists of a cyst of 15 nurse cells and one oocyte enveloped by a single layer of somatic follicle cells (FCs), which constitutes the FE (King, 1970). Oogenesis spans approximately one week and proceeds through 14 developmental stages (S) to eventually give rise to mature eggs (S1 to S14; Fig. 1A) (King, 1970). When the egg chamber buds off from the germarium, approximately 80 FCs enclose the germline cyst. FCs continue to divide mitotically until S6, when they exit the mitotic cycle and switch to an endocycle (Calvi et al., 1998). The apical surface of FCs faces the germline cells, whereas the basal surface contacts the BM, which encapsulates the egg chamber (Gutzeit et al., 1991) (Fig. 1A). Maintenance of FE architecture depends on polarity regulators, cytoskeletal organization, and integrin-mediated adhesion (Abdelilah-Seyfried et al., 2003; Bilder, 2004; Fernández-Miñán et al., 2007; Goode and Perrimon, 1997; Lee et al., 1997). In particular, loss of integrins in the FE results in multilayer formation, highlighting their essential function(s) (Fig. 1B). Two temporally and spatially distinct roles have been proposed: one implicates integrins in regulating mitotic spindle orientation during cell division, paralleling observations in vertebrate epithelia (Fernández-Miñán et al., 2007); the other suggests a role in orchestrating cellular rearrangements during the early stages of FE development (Bergstralh et al., 2013; Lovegrove et al., 2019). Notably, epithelial integrity appears to be particularly vulnerable at the poles, where loss of polarity components or integrins leads to multilayering (Abdelilah-Seyfried et al., 2003; Fernández-Miñán et al., 2007; Goode and Perrimon, 1997). However, the mechanistic basis underlying this regional susceptibility remains elusive.

**Figure 1.**
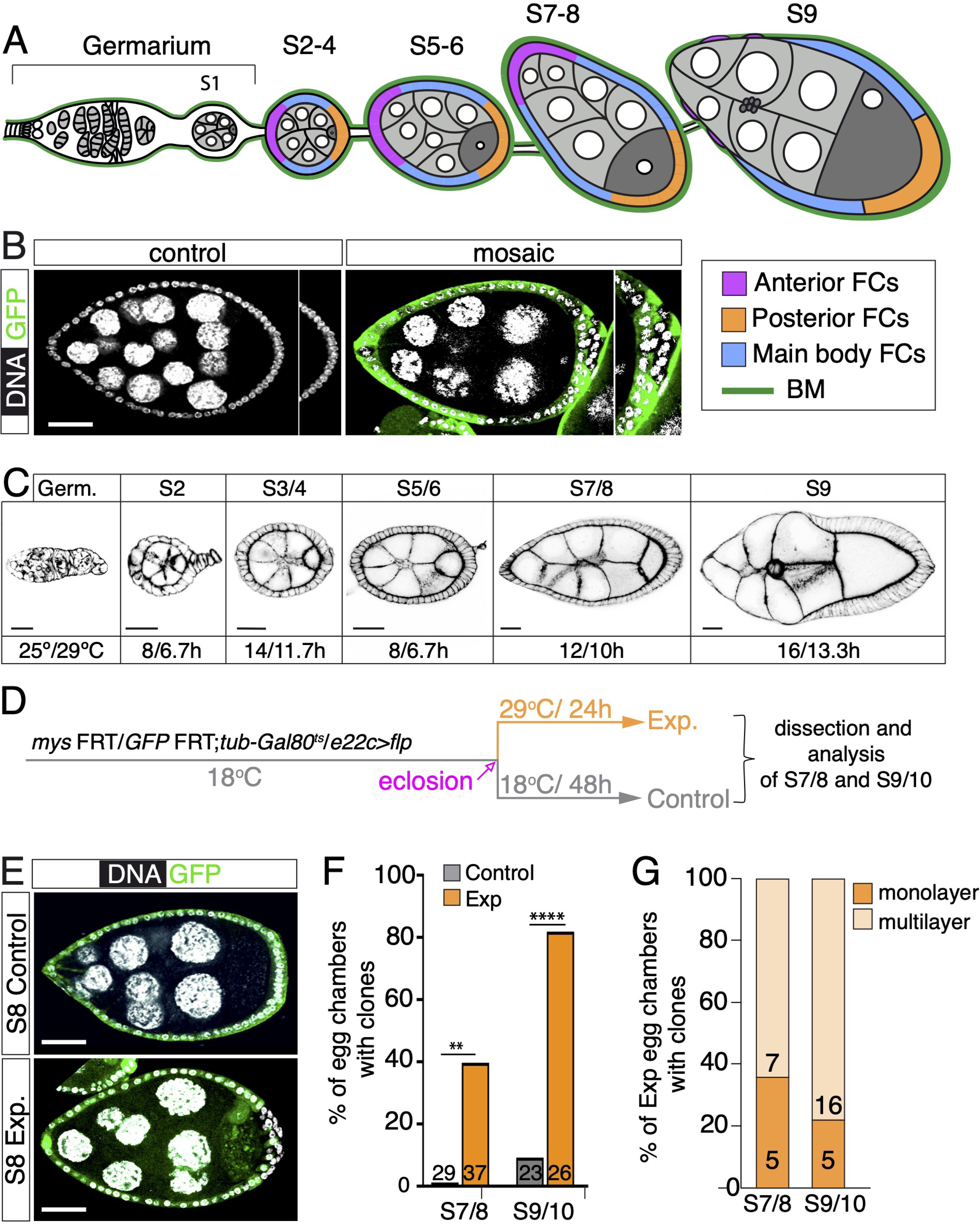
Generation of integrin mutant FCs after the germarium results in multilayering. (A) Schematic representation of a *Drosophila* ovariole showing the germarium and egg chambers up to S9 (S1-S9). (B) S7 control (left) and mosaic (right) egg chambers containing *mys-* FC clones, stained with the DNA marker TO-PRO-3 (white) and anti-GFP (green). Mutant FCs are identified by the absence of GFP. FCs: follicle cells; BM: basement membrane. (C) Control germarium and egg chambers visualized with the F-actin marker rhodamine-phalloidin (black). Developmental timings (in hours) at 25°C and 29°C are indicated for selected stages. (D) Experimental design for clonal analysis. Control flies (always kept at 18°C) were dissected 48h after eclosion, whereas experimental flies were shifted to 29°C for 24h prior to dissection. (E) Control (top) and experimental (bottom) S7 egg chambers stained with TO-PRO-3 to visualize DNA (white) and anti-GFP (green). (F) Quantification of the frequency of egg chambers harbouring *mys* mutant follicle cell clones. (G) Quantification of the multilayering phenotype in mosaic egg chambers carrying *mys* mutant follicle cell clones. Statistical significance was assessed using *Chi square* test, comparing observed and expected frequencies. In this and the rest of figures, P values are the following: * <0.05, ** <0.01, *** <0.001 and **** <0.0001. ns: not significant. Scale bars, 20 μm.

In this work, we address the ongoing debate surrounding integrin function and uncover mechanisms by which integrins maintain simple epithelial organization, as well as those underlying regional susceptibility in the *Drosophila* FE. Combining live imaging, biophysical approaches and quantitative image analysis, we demonstrate that integrin loss induces a stepwise disruption of monolayer organization, culminating in epithelial multilayering. An initial ectopic layer, composed of both control and mutant cells, arises from altered division plane orientation and delayed reintegration following non-planar divisions. Within this layer, exacerbation of these defects in integrin-mutant cells, together with increased cortical tension, synergize to drive and maintain progressive epithelial disruption. In addition, we show that the mechanical properties of the BM contribute to the preservation of simple epithelial architecture. More broadly, our work reveals how integrin-mediated adhesion and BM mechanics coordinate to maintain epithelial architecture, principles likely applicable across diverse epithelial systems in development and disease.

## Results

### Integrins are required to preserve the FE monolayered architecture

The *Drosophila* genome encodes two β integrin subunits—βPS and βν—and five α subunits, designated αPS1 through αPS5 (Brown, 2000; Yee and Hynes, 1993). Of these, βPS, encoded by the *myospheroid* (*mys*) gene, is the only β integrin subunit expressed in the ovary (Brown, 2000; Fernández-Miñán et al., 2007; Yee and Hynes, 1993).

As mentioned above, the role of integrins in the formation and maintenance of the FE remains contested. Our original model proposed that integrins regulated spindle orientation in dividing follicle FCs, whereas other work has suggested that they are required for maintaining cellular organization and morphology only in the germarium (Lovegrove et al., 2019). However, two key methodological differences distinguish our previous study from that of Lovegrove et al. (2019). First, our approach abrogated integrin function throughout FC development, including both early and late stages. Second, while Lovegrove et al. reduced integrin levels in all FCs by expressing a *mys*-targeting RNAi, we employed mitotic recombination to generate mosaic FE containing a mixture of control and *mys* null mutant FCs (*mys-* FCs). To determine whether methodological differences could explain the discrepancies in previously reported phenotypes, here we used a system that allows temporal control of clone induction through temperature. In this system, expression of the recombinase that generates mutant clones is inhibited at 18 °C but activated at 29 °C. Thus, maintaining flies at 18 °C largely prevents clone induction, whereas shifting them to 29 °C triggers clone formation and defines the timing of induction. Based on established egg chamber progression rates at different temperatures (Combedazou et al., 2017; Dillon et al., 2007), stage 7–10 egg chambers raised at 29 °C and analyzed 24 hours after the temperature shift contain *mys-* mutant progeny derived from clones induced at post-germarial stages 3–6 (see Materials and Methods; Fig. 1C-E).

Consistent with repression of clone induction at low temperature, control flies maintained continuously at 18 °C showed *mys-* clones in only 1.2% of stage 7/8 egg chambers and 9.2% of stage 9/10. In contrast, flies shifted to 29 °C displayed a markedly higher frequency of *mys-*clones at the same stages (39.7% and 81.9%, respectively; Fig. 1F), confirming efficient induction after the temperature shift. Detailed analysis further revealed that 64% and 77% of these experimentally induced clones exhibited multilayering, a phenotype never observed in control clones (Fig. 1G).

These findings confirm that integrins are not only essential for maintaining cellular organization and morphology in the germarium (Lovegrove et al., 2019), but also for preserving the monolayered architecture of the FE during its development beyond the germarium. We next sought to investigate the mechanisms through which integrins fulfill this role.

### Division orientation and reintegration differ between poles and main body and depend on integrins

To address the role of integrins in spindle orientation, we performed live imaging of cell divisions in control and *mys-* FCs during S3–6 of egg chamber development. To assess regional differences in multilayer formation, we distinguished between divisions in the main body and in the polar regions, defining the latter as within 3–4 FC diameters of the morphologically distinct polar cells. Because integrin loss does not always cause multilayering and epithelial disorganization may influence division behavior, we analyzed spindle orientation in *mys-* cells within egg chambers that remain monolayered. In addition, this approach allowed us to examine division orientation before multilayering arises and thus helped clarify how multilayering developed over time.

Previous *in vivo* studies showed planar and non-planar FC divisions (Bergstralh et al., 2015), Fig. 2A-C’). To test whether integrin loss altered this behavior, we performed live imaging of S5-6 control and *mys-* FCs (Movies S1, S2). In control egg chambers, non-planar divisions occurred less frequently in the mainbody (17.8%, n=52) than at the poles (30.6%, n=36), revealing an intrinsic regional difference in division orientation within control FE (Fig. 2D, F). In mosaic monolayered egg chambers containing *mys-* clones, the proportion of non-planar divisions among mutant FCs in the main body was significantly higher than in controls (39.4%, n=33 vs 17.8%, n=52). Similarly, non-planar divisions at the poles were more frequent in mutant cells than in controls (60%, n=25 vs 30.6%, n=36; Fig. 2E, F).

**Figure 2.**
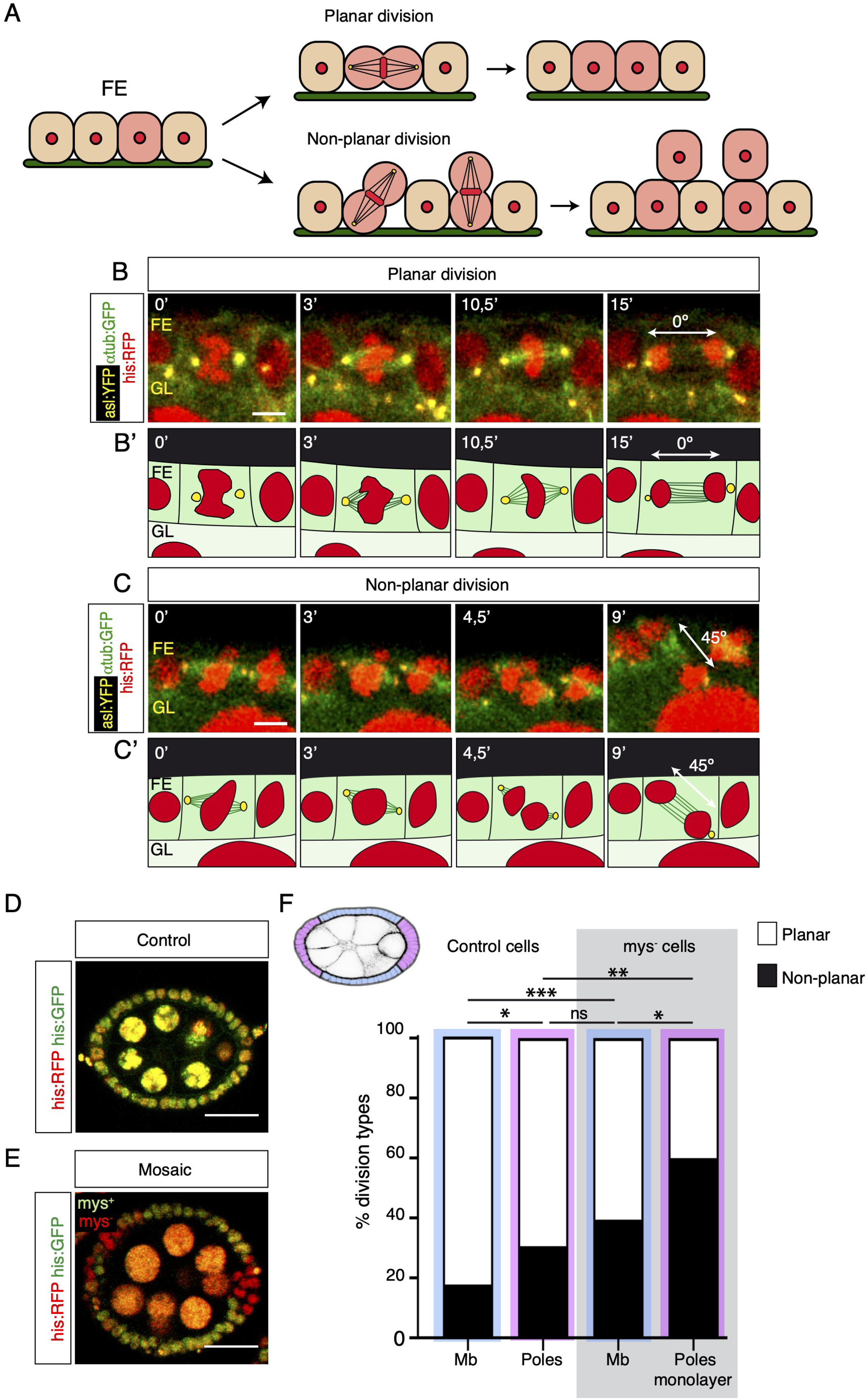
The distribution of planar *vs* non-planar FC divisions depends on both, integrin function and FE subdomain. (A) Schematic representation of planar and non-planar divisions in the FE. (B, C) Confocal images of live S6 egg chambers expressing *histone:RFP* (his:RFP, chromatin, red), *αtubulin:GFP* (αtub:GFP, microtubules, green) and *asterless:YFP* (asl:YFP, centrosomes, yellow). (B, B’) Planar division and (C, C’) non-planar division in the main body. FE, follicular epithelium; GL, germline. (D, E) Confocal images of live egg chambers expressing *histone:RFP* (red) and *histone:GFP* (green). (D) S6 control. (E) S6 mosaic monolayered egg chamber containing *mys-* FC clones in the anterior and posterior terminal domains. Mutant FCs are marked by the absence of his:GFP (green). (F) Quantification of planar and non-planar division frequencies in control and mosaic egg chambers harbouring *mys-* mutant clones, analyzed by regions: main body (Mb) and poles. In this panel and subsequent figures, the schematic in the top left corner depicts a S6 egg chamber, with the main body shaded in pale blue and the poles in pink. Statistical significance was assessed using *Chi square* test, comparing observed and expected frequencies.

Earlier work has shown that divisions occurring at an angle did not necessarily disrupt epithelial architecture, as displaced daughter cells could reintegrate into the monolayer (Bergstralh et al., 2015). To investigate whether integrins play a role in this reintegration process, we conducted live imaging to monitor and characterize reintegration dynamics in both control and mosaic epithelia. In control egg chambers, FCs dividing in the mainbody required significantly less time to reintegrate (mean 16.5 min., n=12) than those at the poles (mean 33 min., n=19) (Fig. 3A, C; Movies S3, S4). In mosaic monolayered egg chambers, *mys-* FCs in the main body exhibited prolonged reintegration times (mean 44.3 min., n=16) relative to controls (Fig. 3B, C; Movie S5). Similarly, *mys-* FCs at the poles required significantly more time (mean 65.57 min., n=14) to reintegrate than both control FCs at the poles and mutant cells in the main body (Fig. 3C; Movie S6).

**Figure 3.**
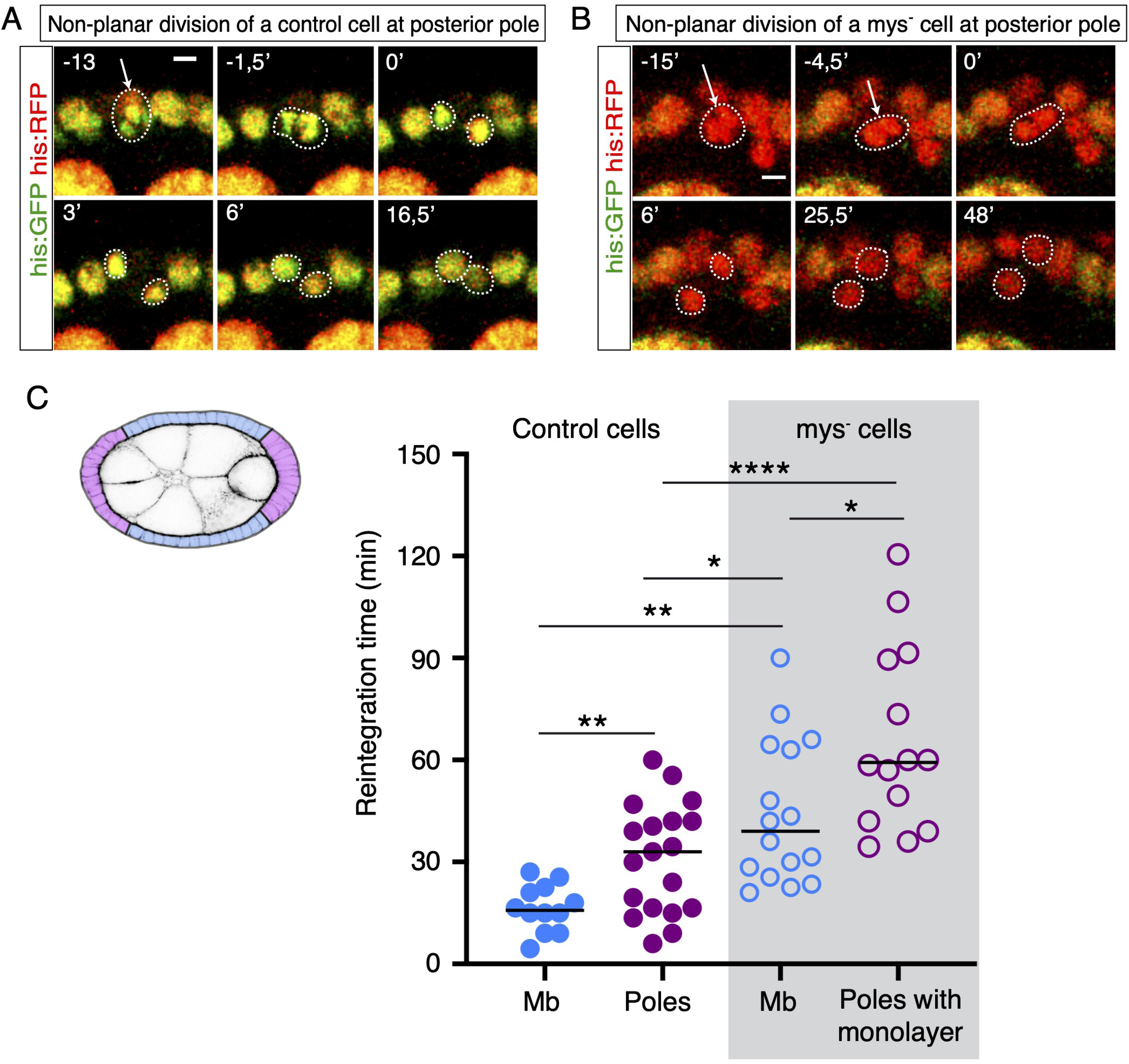
Reintegration times following non-planar divisions in FE monolayers depend on integrin function and position within the egg chamber (main body vs poles). (A, B) Confocal images of live S5 egg chambers expressing *histone:RFP* (his:RFP, red) and *histone:GFP* (his:GFP, green) showing cell reintegration dynamics following non-planar divisions at the poles of control (A) and mosaic monolayered egg chambers containing *mys-*FC clones, identified by the absence of his:GFP (B). (C) Quantification of reintegration times in control and mosaic egg chambers harbouring *mys-* clones, analyzed by region: main body (Mb) and poles. Statistical significance was assessed using a Student’s *t-test*. Error bars indicate standard error of the mean (SEM). Scale bars, 3 μm.

Altogether, these results indicate that integrins are essential not only for controlling division orientation, but also for ensuring efficient reintegration of daughter cells upon non-planar FC divisions, thereby preserving epithelial architecture. Our data further suggest the existence of a threshold reintegration time beyond which monolayer integrity becomes compromised. The prolonged reintegration of integrin mutant cells at the poles may therefore explain the regional specificity of the multilayering phenotype. Indeed, extended residence outside the epithelial layer increases the likelihood that displaced cells divide again before reintegrating, contributing to multilayer formation. Moreover, once cells occupy ectopic positions, their behavior may be further influenced by the disorganized environment, potentially exacerbating these defects. To examine this possibility, we analyzed the behavior of integrin mutant cells in mosaic egg chambers displaying a multilayered follicular epithelium.

### Integrin-deficient cells in ectopic layers display enhanced defects that reinforce multilayering

To test whether integrin-deficient cells within ectopic layers contribute to multilayering at the poles, we compared the behavior of *mys-*FCs in multilayered versus monolayered mutant FE, as well as to FCs in control FE (Fig.4A-C). Non-planar divisions were more frequent in *mys-*FCs within ectopic layers than in mutant cells in monolayered FE (71.5%, n=35 vs 52%, n=29; Fig. 4A) or in control FCs (28.2%, n=32; Fig. 4D), indicating that division defects are enhanced in ectopic layers. Similarly, we found that *mys-* FCs located in ectopic layers exhibited a significantly longer reintegration time (mean 108.23 min., n=17; Movie S7) than mutant cells in a monolayered FE (mean 61,3 min., n =15; Movie S6) or control FCs (mean 31.13 min, n=19; Fig. 4E; Movie S4). Of interest, 29.4% of *mys-* FCs in the ectopic layers fail to reintegrate during the time window of the live imaging (triangles in Fig. 4E). Last, the ectopic layers in mosaic egg chambers contained both integrin-positive and integrin-negative cells. To determine whether the enhanced defects of integrin mutant cells in multilayered epithelia reflect integrin loss rather than simply a disorganized environment, we compared heterozygous *mys*+/- FCs with homozygous *mys-* FCs residing within ectopic layers. We found that *mys*+/-FCs divided non-planarly at frequencies comparable to controls (34.2%, n=41) and significantly lower than *mys*-FCs (71.4%, n=35; Fig. 4D). Furthermore, the time required for *mys*+/- cells in ectopic layers to reintegrate (mean 40.4 min., n=16) was not significantly different from that of controls (mean 31.13 min., n=19; Fig. 4E). These results indicate that the increased frequency of non-planar divisions and the prolonged reintegration time observed in ectopic layers are specifically caused by loss of integrin function. Together, they suggest that once integrin-deficient cells become displaced into an ectopic layer, their defects are further exacerbated, promoting additional perpendicular divisions and delayed reintegration. Hence, this positive feedback likely reinforces progressive epithelial disorganization.

**Figure 4.**
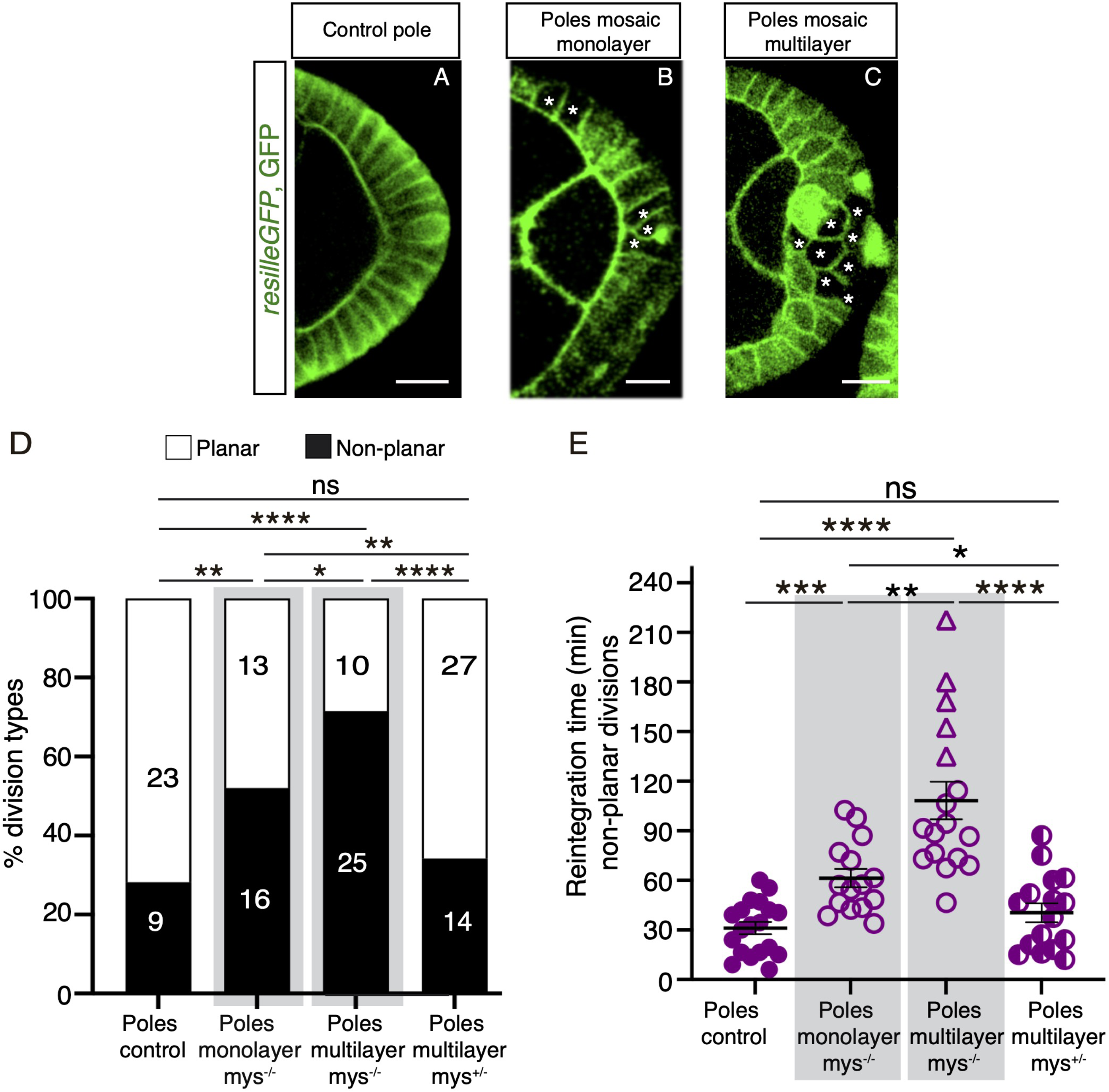
Integrins regulate division plane and reintegration time in an architecture-dependent epithelial context. (A-C) Posterior pole regions of S5 control (A), monolayered (B) and multilayered (C) mosaic egg chamber expressing *resille:GFP* and containing *mys-* FC clones, identified by the absence of his:GFP. (D) Quantification of division orientation frequencies and (E) reintegration times at the poles in controls and mosaic egg chambers carrying *mys-* and *mys*+/- FCs, with or without multilayering. Triangles indicate FCs that fail to reintegrate during imaging time window. Statistical significance was assessed using a *Chi square* test, comparing observed and expected frequencies (A) and a Student’s *t-test* (B). Error bars indicate SEM. Scale bar, 3 μm.

### Integrin regulation of junctional tension contributes to epithelial monolayer disruption

Previous studies have shown that local increases in cortical actomyosin contractility at cell-cell interfaces can promote cell sorting (Brodland, 2002). We previously reported that loss of integrin function leads to increased junctional tension at cell-cell contacts in S9 FCs (Santa-Cruz Mateos et al., 2020). In this context, elevated surface tension between integrin mutant cells within ectopic layers could promote their separation from the primary layer, reducing the likelihood of reintegration, thereby facilitating the stabilization and expansion of multilayering. Consistent with this idea, we previously showed that increasing cell tension enhances the multilayering phenotype associated with integrin loss of function (Ng et al., 2016). To explore this hypothesis, we quantified junctional tension at cell-cell contacts in control and *mys-* FCs.

Laser ablation at cell-cell contacts is a well-established method to measure junctional tension (Farhadifar et al., 2007). We therefore performed laser ablation experiments on the basal side of S5–6 control and *mys-* FCs using a UV laser. Membrane dynamics were visualized using Resille:GFP (Morin et al., 2001). For simplicity, we assumed equivalent cytoplasmic viscosity in *mys-* and control FCs. To minimize potential effects of anisotropic forces within the follicular epithelium (FE), all cuts were made parallel to the anterior–posterior axis. Ablations were performed in both the polar and main body regions to assess potential regional differences (Fig. 5A, B). As expected, laser ablation caused junctional relaxation, increasing the distance between cell vertices flanking the ablation site (Fig. 5B; Movie S8). In control egg chambers, the initial retraction velocity was similar in the main body (0.43 µm/s, n=16) and pole regions (0.42 µm/s, n=27). In addition, *mys-* FCs in monolayered FE showed comparable velocities (0.40 µm/s in the main body, n=19; 0.42 µm/s at the poles, n=22). In contrast, *mys-*FCs within multilayered regions displayed significantly higher vertex displacement, with an average retraction velocity of 0.65 µm/s, (n=19; Fig. 5C).

**Figure 5.**
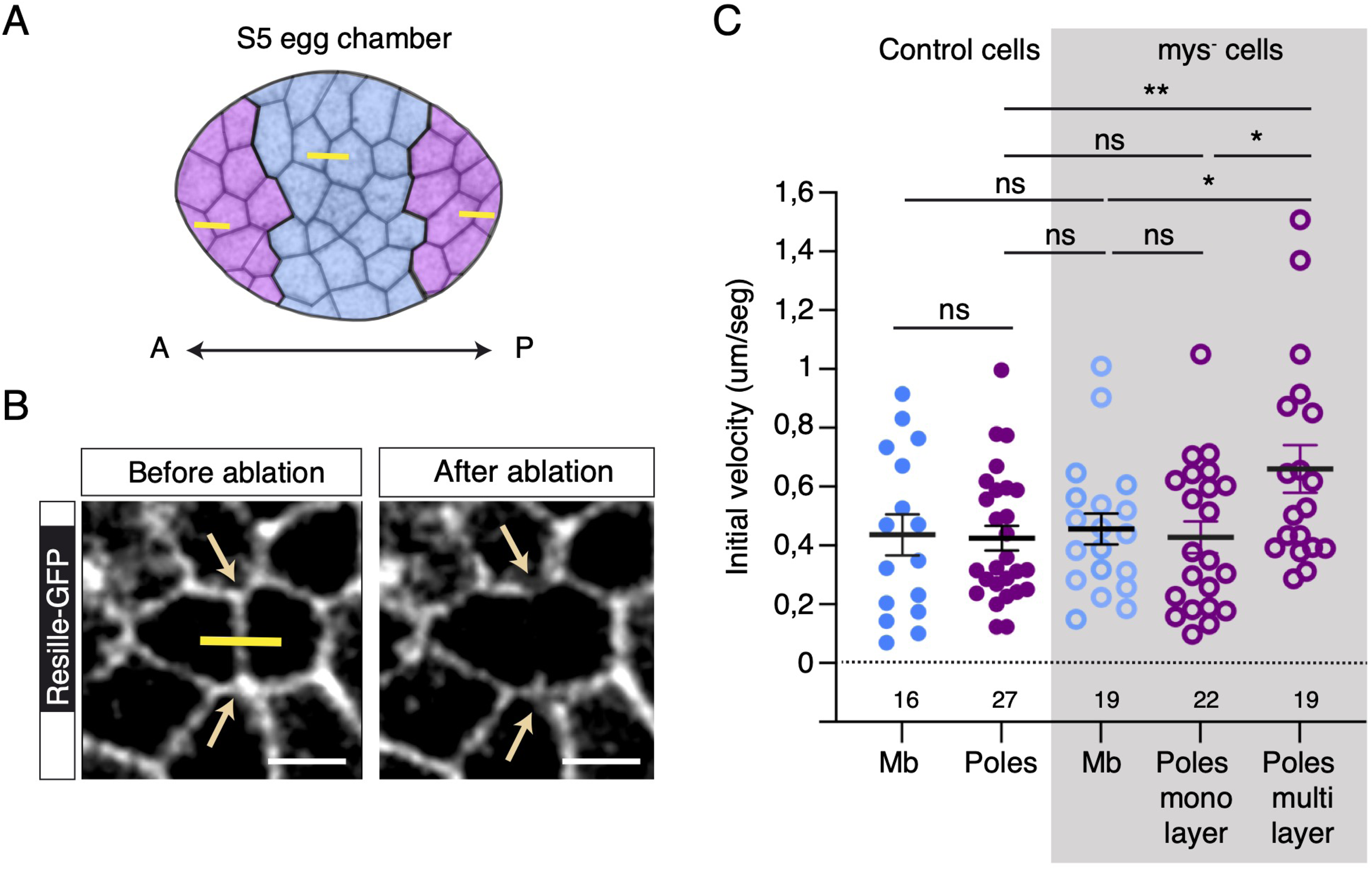
Integrins regulate junctional tension in an architecture-dependent epithelial context. (A) Schematic of an S5 egg chamber, with the main body region shaded in pale blue and the poles in pink, indicating the orientation and position of laser ablations. All ablations were performed parallel to the AP axis (A: anterior; P: posterior). (B) Representative images of a live S5 control egg chamber expressing *resille:GFP*, before and after laser ablation of an individual cell-cell junction. (C) Quantification of initial vertex displacement velocity following ablation of the indicated cell junctions. Statistical significance was assessed using a Student’s *t-test*. Error bars indicate SEM. Scale bar, 3 μm.

Together, these results support a multistep model in which integrin loss triggers a reinforcing cascade of defects (aberrant division orientation, delayed reintegration, and increased junctional tension), which synergize to progressively drive multilayering. As these defects accumulate over time, this model predicts that the severity of the multilayering phenotype should increase as egg chamber development advances. Consistent with this prediction, we observed a stage-dependent rise in the proportion of mosaic egg chambers displaying a multilayered phenotype (Supp. Fig. 1).

### BM mechanics as a key factor in maintaining simple FE integrity

Measurements of BM stiffness in the *Drosophila* egg chambers have revealed a mechanical anisotropy in the BM, with the poles being approximately 50% softer than the central regions from early S3 (Crest et al., 2017). In our study, we observed a clear asymmetry in both division plane orientation and reintegration time of FCs between the poles and the main body of control egg chambers (Figs. 2 and 3). Moreover, loss of integrins led to multilayer formation specifically at the poles, even though mutant follicle cells in the main body also showed increased non-planar divisions and delayed reintegration. These observations suggest that regional differences in BM stiffness could contribute to the observed phenotypic differences. To test whether BM stiffness heterogeneity contributes to multilayering in the absence of integrins, we analyzed the behavior of integrin mutant cells in egg chambers with homogenously softened BMs. Egg chambers uniformly depleted of laminins or Collagen IV (Col IV) exhibit homogenously soft BMs (Diaz de la Loza et al., 2017). We therefore generated *mys* mutant clones in egg chambers depleted of either laminins or ColIV. Laminin levels were reduced in all FCs, using the *tj*-*Gal4* driver to express an RNAi construct targeting *Laminin B1* (*LanB1*), which encodes the sole β chain shared by the two Laminin trimers present in *Drosophila* (Diaz de la Loza et al., 2017; Lee et al., 2003). Similarly, Col IV levels were reduced using *tj*-*Gal4* to drive to express an RNAi construct against *viking* (*vkg*), which encodes the ColIVα2 subunits (*tj*>*vkg* RNAi; Supp. Fig. S2A-F). Strikingly, reducing either Laminin or ColIV levels substantially rescued the multilayering phenotype. While 78,37% of mosaic egg chambers containing *mys-* clones exhibited multilayering (n=21), only 26% of *tj*>*LanB1* RNAi (n=58) and 28,57% of *tj*>*vkg* RNAi egg chambers with *mys* clones (n=56) developed a stratified epithelium (Supp. Fig. 2G).

Together, these findings support a model in which BM stiffness heterogeneity contributes to the disruption of monolayered architecture in the absence of integrins. We next asked which cellular behaviors—such as division orientation or reintegration time—mediated this effect. While mosaic clones in laminin- or Col IV–depleted egg chambers can be generated, scaling this up to perform sufficient *in vivo* experiments for statistical analysis is technically very challenging. We therefore turned to an alternative approach. Because integrin downregulation in the FE reduces laminin levels, global integrin depletion is expected to produce a uniformly softened BM, providing an ideal context to test the role of BM stiffness heterogeneity in multilayer formation in the absence of integrins. To reduce integrin levels throughout the FE, we expressed an RNAi construct targeting *mys* under the control of the *tj*-Gal4 driver (*tj>mys RNAi*,) and measured BM stiffness. Although previous studies have shown that strong integrin knockdown can result in severe disorganization of the ovariole structure (Bergstralh et al., 2013), thereby complicating the analysis of FE-specific phenotypes, we used an alternative *mys RNAi* line (Martin-Bermudo and Brown, 1999; Valencia-Exposito et al., 2022), that reduces integrin levels without inducing major structural defects (Fig. 6A, B). In this case, multilayering in S7-8 egg chambers was observed in a fraction of FE (15%, n = 40; Fig. 6C), compared with mosaic egg chambers containing *mys-* clones (78,37%, n=21; Sup. Fig. 2G). Moreover, when present, multilayering remained limited and never exceeded two layers. Atomic force microscopy measurements confirmed that *mys*-depleted follicles exhibited significantly reduced BM stiffness compared to controls and, importantly, lacked the typical anisotropic stiffness distribution observed in wild-type egg chambers (Fig. 6D).

**Figure 6.**
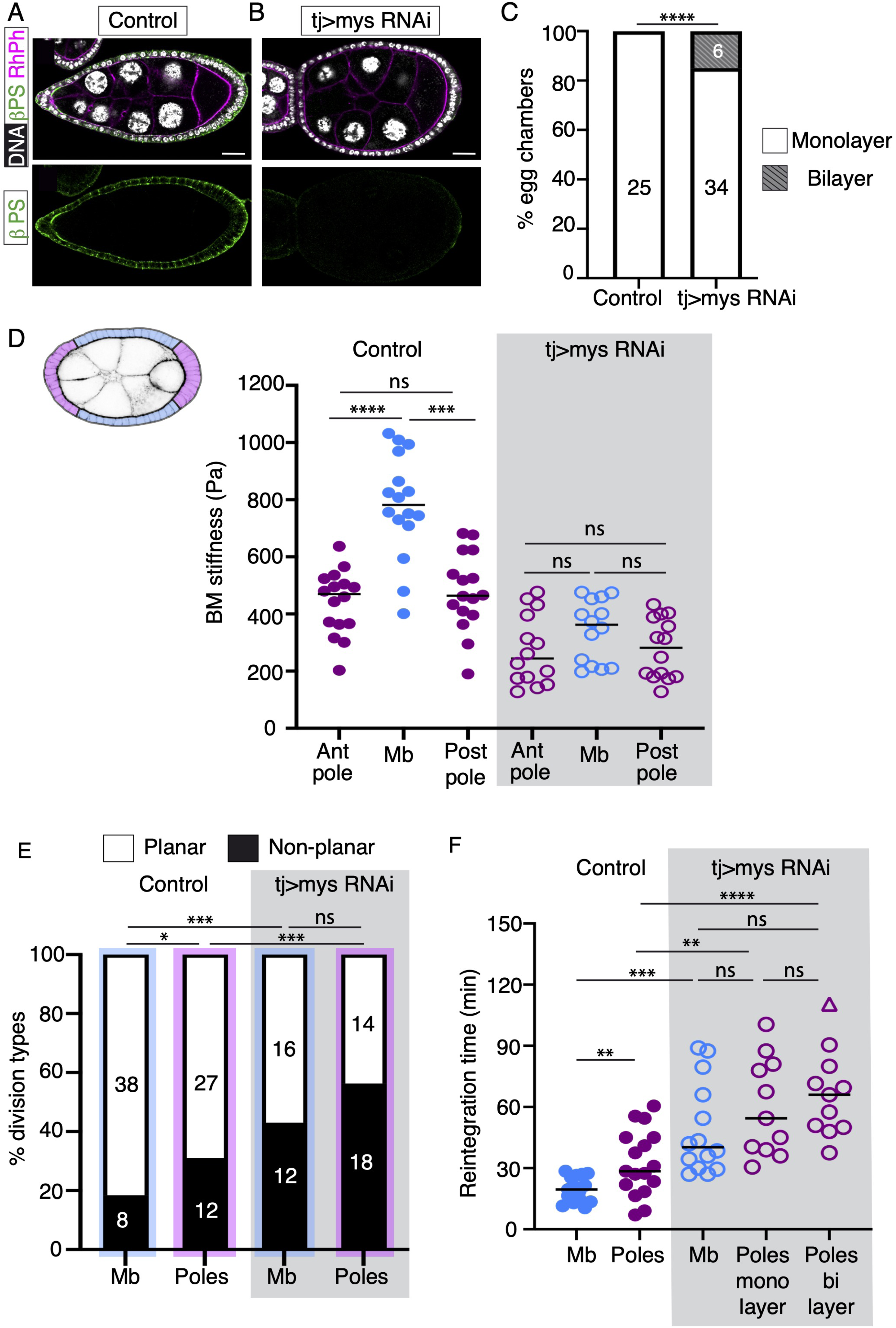
Basement membrane stiffness anisotropy influences epithelial multilayering. (A) Control egg chamber and (B) experimental egg chamber with uniform *mys* depletion (*tj>mys RNAi*) stained to visualize DNA (white, top), F-actin (magenta, top) and the βPS integrin subunit (green, top and bottom). (C) Quantification of the multilayering phenotype in control and *tj>mys RNAi* egg chambers. (D) Comparison of the apparent elastic modulus (*K*) in the main body (Mb), anterior (Ant) and posterior (Post) pole regions of control and *tj>mys RNAi* egg chambers. (E, F) Quantification of planar vs non-planar division frequencies (E) and reintegration times (F) in control and *tj>mys* RNAi egg chambers. The triangle indicates a FC that failed to reintegrate during imaging. Statistical significance was assessed using a Student’s *t-test*. Error bars indicate SEM. Scale bar, 20 μm (A, B).

We then examined division orientation and reintegration dynamics by live imaging using the membrane marker Resille:GFP (Movie S9, S10). In control egg chambers, non-planar divisions showed regional differences, occurring in 18.4% of cells in the main body and 31% at the poles (Fig. 6E). In contrast, *tj-Resille:GFP>mys* RNAi egg chambers showed no regional bias, with similarly elevated frequencies in both regions (43% in the main body and 56.3% at the poles). Reintegration dynamics followed a similar trend. In controls, reintegration was slower at the poles (27.5 min., n=17) than in the main body (19.5 min., n=17). In *tj-Resille:GFP>mys RNAi* egg chambers, reintegration was markedly delayed across the entire FE, supporting a role for integrins in regulating reintegration timing. However, no significant regional differences were observed in this condition, with reintegration times of 40.25 min. in the main body (n=14), and 54.5 min. (n=11) and 66 min. (n=11) at the poles of monolayered and bilayered epithelia, respectively (Fig. 6F). Notably, reintegration times in *tj-Resille:GFP>mys* RNAi egg chambers were shorter than those observed for *mys* mutant FCs in mosaic egg chambers, which retain BM stiffness anisotropy (mean: 61.3 min. and 108.23 min. in monolayered and multilayered FE, respectively). This difference may explain the lower frequency of multilayering in *tj-Resille:GFP>mys* RNAi egg chambers compared to mosaics. Nevertheless, reintegration time tended to increase in the main body of mosaic monolayered with respect to bilayered *tj-Resille:GFP>mys RNAi* egg chambers. Moreover, the mean reintegration time in the latter (66 min.) slightly exceeded that of monolayered mosaic egg chambers (61.3 min.). Together with the observation of a *mys* RNAi FC that failed to reintegrate within the imaging period (110.5 min.; triangle in Fig. 6F), these findings may explain the rare occurrence of bilayer formation in *tj-Resille:GFP>mys RNAi* samples, while overall supporting the idea that BM stiffness heterogeneity contributes to the disruption of monolayered architecture when integrin function is compromised.

These results suggest that integrins directly regulate division orientation, whereas their influence on reintegration could be modulated by regional BM mechanical properties. They further indicate that regional differences in BM stiffness amplify the impact of integrin loss on reintegration, contributing to the disruption of monolayered architecture.

## Discussion

Simple epithelia form continuous cell layers supported by a basement membrane and lining internal and external body surfaces. Although they are essential for tissue function, the mechanisms that maintain their architecture remain less well understood. Here, we used the Drosophila follicular epithelium as a model to investigate how simple epithelial integrity is preserved. Our findings show that integrins preserve monolayer organization by preventing a progressive disruption that culminates in epithelial multilayering, in a context modulated by BM mechanics. Mechanistically, this function relies on controlling division plane orientation, timely reintegration after non-planar divisions, and junctional tension, which together safeguard epithelial integrity.

Previous studies support a role for integrins in maintaining simple epithelial architecture and preventing stratification. Loss of Itgb1 in mice induces multilayering in skin and lung epithelia, and its downregulation is part of normal epidermal stratification (Brakebusch et al., 2000; Jones and Watt, 1993; Simpson et al., 2011). Our findings extend this framework by defining the cellular mechanisms through which integrins preserve the monolayered FE in *Drosophila*. Consistent with earlier work (Fernández-Miñán et al., 2007), integrin-deficient FCs show a bias toward non-planar divisions. This bias is further enhanced in ectopic layers, where continued non-planar divisions likely reinforce multilayering, in line with observations in mouse skin where suprabasal divisions contribute to stratification (Damen et al., 2021). Together, these results strengthen the link between integrins and mitotic spindle orientation. In cell culture systems, integrin β1 has been proposed to regulate spindle positioning via a ligand-independent mechanosensory complex, comprising FAK, Src, and p130Cas (Petridou and Skourides, 2016). Future work should determine whether this pathway is conserved *in vivo* and operates within intact epithelial tissues, as well as elucidate the molecular mechanisms by which integrins regulate cell reintegration dynamics.

Although our findings support a role for integrins in controlling spindle orientation and the plane of division (Fernández-Miñán et al., 2007; Petridou and Skourides, 2016), a compelling aspect of multilayer formation in the *Drosophila* FE is that integrin depletion does not invariably lead to multilayering. This indicates that altered division orientation alone is insufficient to disrupt epithelial architecture. In fact, while numerous studies in mammalian epithelia, including the skin, hair follicle, tongue and prostate, have highlighted the importance of oriented cell divisions in ensuring proper morphogenesis and structure of epithelial tissues (Byrd et al., 2016; Lechler and Fuchs, 2005; Schaeffer et al., 2004; Seldin et al., 2016), other findings challenge this view. For example, skin explant cultures treated with inhibitors of cell division retain some capacity to stratify and differentiate (Damen et al., 2021). In addition, in embryonic mouse kidney, zebrafish neuroepithelial cells, and multiple *Drosophila* tissues, non-planar divisions are tolerated, with displaced cells reintegrating into the epithelial layer to preserve tissue integrity (Bergstralh et al., 2015; Ciruna et al., 2006; Packard et al., 2013). Consistent with these observations, our data identify reintegration timing following non-planar divisions, a process also regulated by integrin function, as a critical determinant of simple epithelial integrity. Notably, in multilayered egg chambers, mutant cells exhibit prolonged reintegration times compared to both control and mutant cells in non-multilayered egg chambers. These findings suggest that epithelial disruption arises when reintegration is delayed beyond a critical threshold. Such delays likely increase the probability that displaced cells divide outside the monolayer, thereby promoting the accumulation of ectopic layers. Our results further reveal that integrin mutant cells within ectopic layers display elevated junctional tension relative to both control cells and mutant cells in monolayers. Increased tension may additionally stabilize ectopic positioning by limiting reintegration, consistent with previous findings indicating that enhanced cortical tension exacerbates multilayering upon integrin loss (Ng et al., 2016). This is also in agreement with *in vitro* studies of epidermal stratification, where delaminated cells adopt a high-tension state that reinforces separation between basal and suprabasal layers (Miroshnikova et al., 2018). Together, our results support a multifactorial model of epithelial disruption in which division orientation, reintegration dynamics, and junctional tension act in concert (Fig. 7). Integrins coordinate these processes to preserve monolayer integrity; their loss shifts this balance, enabling the progressive formation and stabilization of ectopic layers.

**Figure 7.**
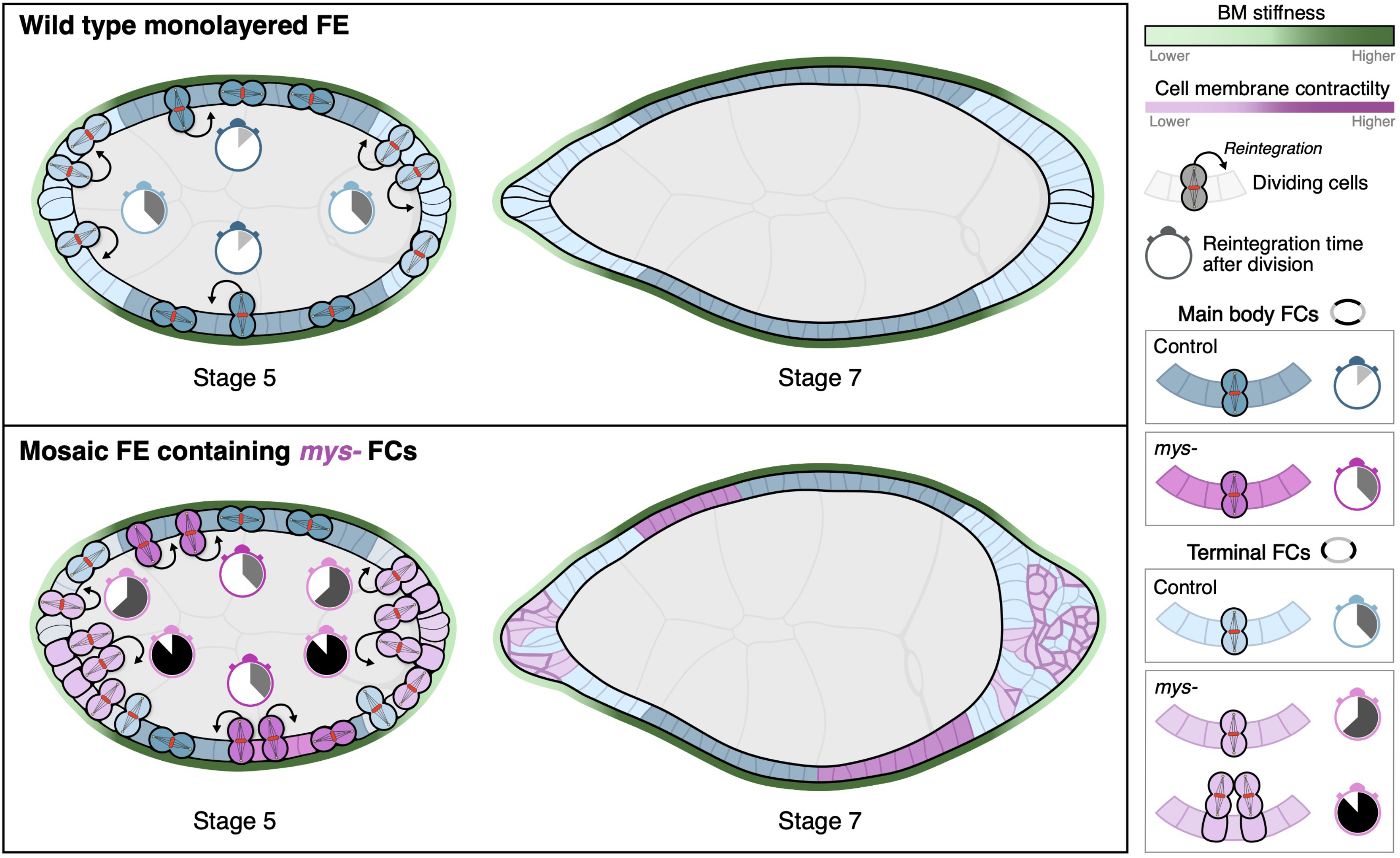
Model illustrating the mechanisms by which integrins and BM stiffness align to safeguard simple epithelial architecture. Drawings of S5 and S7 wild-type (top) and mosaic egg chambers containing *mys^-^* FCs (bottom). The FE is divided into poles (light blue) and main body (dark blue) regions. Up to S6, FCs divide either parallel to the epithelial plane (planar divisions) or at an angle (non-planar divisions). In wild-type epithelia, non-planar divisions occur more frequent at the poles; however, epithelial architecture is preserved through reintegration, which is slower at the poles (chronometers). In mosaic egg chambers, *mys^-^* FCs exhibit higher frequencies of non-planar divisions and delayed reintegration compared to *mys^+/-^* cells, particularly at the poles and in mutant cells in extra layers. These delays, together with reduced BM stiffness at the poles (green) and increased contractility of *mys^-^* FCs in extra layers (stronger pink lines), impair reintegration, ultimately disrupting epithelial organization and promoting multilayer formation.

Multilayering of the FE has been reported in mutants affecting apico-basal polarity, cytoskeletal organization and Hippo signaling (Meignin et al., 2007; Ng et al., 2016; Polesello and Tapon, 2007; Tanentzapf et al., 2000). Intriguingly, in all cases, multilayering is restricted to the poles. The basis for this spatial specificity remains unclear. Our findings point to regional differences in BM mechanics as a key determinant. Supporting this idea, a stiff BM has been shown to promote planar-oriented divisions in skeletal muscle stem cells (Moyle et al., 2020). Extending this concept in vivo, we find that BM stiffness correlates with both division orientation and reintegration dynamics in the developing epithelium: stiffer regions, such as the main body, display more frequent planar divisions and faster reintegration, whereas softer regions at the poles show increased non-planar divisions and delayed reintegration. These properties likely render the poles more susceptible to disruption upon loss of factors regulating division orientation. However, integrin loss in a uniformly soft BM does not induce multilayering, indicating that BM softness alone is insufficient. Thus, we propose that spatial heterogeneity in BM stiffness—rather than absolute stiffness—is critical for driving localized epithelial disruption. Such mechanical heterogeneity may enable regional transitions from monolayered to multilayered states, potentially through localized BM remodeling. Consistent with this idea, mammary gland branching involves a transient multilayered intermediate at the leading edge of elongating ducts, in regions lacking laminins (Ewald et al., 2008).

Tissue architecture emerges from the interplay between cellular behaviors and molecular and mechanical cues. Our study reveals that integrins, together with BM mechanics, maintain simple epithelial organization. Disruption of the monolayer appears to be a multistep process: the formation of ectopic layers creates a disorganized environment that further amplifies epithelial disorganization. Such progressive breakdown of tissue architecture is a hallmark of oncogenic transformation, and forces from the underlying basement membrane can influence tumor architecture and malignant potential (Fiore et al., 2020). Understanding the factors that preserve epithelial integrity and regulate transitions to multilayered states is therefore critical for both morphogenesis and cancer progression.

## Materials and methods

### *Drosophila* fly stocks and genetics

The following fly lines were used: *mys^11^* (also known as *mys^XG43^* (Bunch and Brower, 1992)), *e22c-Gal4-UAS-flipase* (Duffy et al., 1998), *tub-Gal80^ts^* (Bloomington *Drosophila* Stock Centre (BDSC) number 7019), *his:RFP* (BDSC 25377), *UAS-mys RNAi* (NIG-Fly, P{NIG.1560R}2), *UAS-LanB1 RNAi* (Vienna *Drosophila* Resource Centre (VDRC) 23121), *UAS-ColIVα*2 *RNAi* (VDRC v106812), *pUbi-YFP:asl*, *pUbi-GFP:αtub 84B* (a gift from Dr. Cayetano González), *resille:GFP* (Morin et al., 2001), *traffic jam (tj)*-*Gal4* (Li et al., 2003), *his2Av:GFP hs-FLP FRT-19A* (BDSC 32045), *y w pUbi*-*GFP FRT-101* (BDSC 5153) and *hs-flp, pUbi-RFP* FRT-19A (BDSC 31418).

To generate *mys* mutant clones, we used either the *e22c-Gal4* driver, which is expressed in somatic cells of the germarium in pupal and adult ovaries, or the heat shock-driven (*hs-flp*) system to induce *flipase* expression. We utilized the *pUbi-GFP FRT-101* to label the clones.

To generate *mys^11^* clones in post-germarial epithelia, *mys^11^ FRT101*/*y w pUbi-GFP FRT-101*; *tub-Gal80^ts^*/*e22c-Gal4 UAS-flp* females were grown at 18°C. Upon eclosion, adults were kept either at 18°C for 48 hours (control condition) or shifted to 29°C for 24 hours (experimental condition). Since progressing from stage 2 to stage 8 takes approximately 40 hours at 25°C (Spradling, 1993), and since *Drosophila* development at 29°C is accelerated by ∼1.2-fold (Dillon et al., 2007), S7/8 and S9/10 clones analyzed 24 hours after the temperature shift to 29°C were induced from stage 2-onwards.

To study wild type FC divisions *in vivo,* we used females carrying *pUbi-YFP:asl*, *pUbi-GFP:αtub*; *his:RFP*. To study *mys^11^*mutant FC divisions *in vivo*, we obtained different types of females: *pUbi-YFP:asl*, *mys^11^* FRT-19A/*his:2AV:GFP hs-flp FRT-19A*; *resille:GFP*, *his:RFP*/*e22c*-*Gal4 UAS*-*flp* and *mys^11^* FRT-19A/*his:2AV:GFP hs-flp FRT-19A*; *his:RFP*/*e22c*-*Gal4 UAS*-*flp* (to generate small clones to analyze cell behaviors in monolayers), and *mys^11^ FRT-101/y w pUbi*-*GFP FRT-101*; *resille:GFP/e22c-Gal4 UAS-flp* (to obtain clones in multilayers and for laser ablation experiments). In these experiments, flies were grown always at 25°C.

For the rescue experiments, we used the heat shock flipase (hs-flp) system (Chou and Perrimon, 1992) to generate follicle cell mutant clones and *tj-Gal4* to express either *UAS*-*LanB1 RNAi* or *UAS*-*ColIVα*2 *RNAi*. Females *mys^11^* FRT-19A;; *UAS*-*LanB1 RNAi* or *mys^11^*FRT-19A;; *UAS*-*ColIVα*2 *RNAi* were crossed to hs flpRFPFRT19A; *tj-Gal4* males. The heat shock was performed at 37 °C for 2 h during third instar larvae and newly hatched females. Flies were kept at 25°C and yeasted for 2 days prior to ovary dissection.

### Ex vivo culturing of egg chambers

For live imaging, 1-2 days-old females were fattened on yeast during 2-3 days before dissection. We follow the protocol used by Villa-Fombuena *et al*., 2021. Briefly, the day before the experiment, a drop of 3 μl of Cell-Tak (Corning 354240) mixed with 3 μl of NaCOH_3_ was added a MatTek and kept at 4°C. Adult ovaries were dissected at room temperature in Ringer’s medium [128 mM NaCl, 2 mM KCl, 1.8 mM CaCl_2_, 4 mM MgCl_2_, 35.5 mM sucrose, 5 mM HEPES (pH 6.9)]. Ovarioles without the muscle sheath were transferred to plate and the plate was then supplemented with Schneider’s medium (Sigma Aldrich). Samples were imaged for a maximum of 4 hours under a scanning confocal microscope.

### Immunohistochemistry

1-2 days-old females were grown 2-3 days at 25°C before dissection. Adult ovaries were dissected at room temperature (RT) in Schneider’s medium (Sigma Aldrich), fixed in 4% paraformaldehyde in PBS (ChemCruz) for 20 min, permeabilized 30 min in PBT and blocked 1 h in PBT-10. Incubation with primary antibodies was performed overnight at 4°C in PBT-1. The following primary antibodies were used: chicken anti-GFP (1:600, Abcam Cat#13970), anti-GFP booster (1:500, Chromotek Cat#17303343), rat anti-RFP (1:500, Chromotek Cat#17812387). Secondary antibodies were incubated for 2 h in PBT-0.1 and used 1:200: anti-chicken Alexa Fluor 488 (Invitrogen Cat# A11039), anti-Rat Texas red (Sigma Aldrich Cat#SAB3700551). To label DNA, ovaries were incubated for 10 min with Hoechst (Sigma-Aldrich, 5 mg/ml; 1:1000 in PBT). For filamentous actin labelling, ovaries were incubated with Rhodamine Phalloidin (Biotium, 1:40) or 488 Phalloidin (Biotium, 1:40), for 15 minutes. Ovaries were mounted in Vectashield (Vector Laboratories).

PBT-10: PBS, 10% BSA, 1% tween20.

PBT-1: PBS, 1% BSA, 1% tween20.

PBT-0.1: PBS, 0.1% BSA, 1% tween20.

PBT: PBS (phosphate-buffered saline), 1% tween20.

### Laser ablation

Laser ablation experiments were performed in an Olympus IX-81 inverted microscope equipped with a spinning disk confocal unit (Yokogawa CSU-X1), a 100× oil objective, a 355 nm pulsed, third-harmonic, solid-state UV laser, and an Evolve 512 EMCCD digital camera (Photometrics). To analyze tension between FC-FC, a pulse of 125 mJ energy and 4 msec. duration was applied to sever plasma membranes of cells. In all cases, cell surfaces were visualized with the membrane marker *resille:GFP* and a Cobolt Calypso state laser (l = 491 nm 50 mW) was used for excitation of the GFP. Cuts between FC-FC were made parallel to the antero-posterior axis of the egg chamber. Images were taken, 3 sec before and 10 sec after laser pulse, every 0,5 sec. The initial velocity was estimated as the velocity at the first time point after ablation (t1= 0,5 sec).

### Atomic force microscopy

Atomic force microscopy measurements were performed as in (Diaz de la Loza et al., 2017; Molina Lopez et al., 2023). Ovarioles were dissected out of the muscle sheath to make sure that the AFM cantilever was in direct contact with the BM. The cantilever was positioned at the desired position by brightfield microscopy. Statistical significance between experimental and control values was evaluated using a t-test, **** and *** P value < 0.00001 and <0.0001, respectively. Sixteen measurements were collected from each region (anterior, main body and posterior) across 6 control egg chambers, and 14 measurements per region from 6 *mys*-depleted egg chambers.

### Imaging and processing of samples

Live sample imaging was acquired on a Leica TCS-SP5 confocal microscope with a 40x/1,3 oil objective and a Zeiss LSM 880 confocal microscope with a 40x/1,2 water objective. In all cases, Z-stacks were taken with 1 μm intervals and time points recorded every 1,5 minutes, for 2-4 hours.

Images from fixed sample were taken with a Leica Stellaris confocal microscope equipped with a 40x/1,3 oil immersion objective. Z-stacks were 1 μm intervals. Both fixed and live images were analyzed utilizing ImageJ and processed with Adobe Photoshop and Adobe Illustrator.

## Statistical analysis

Statistical analysis of categorical data was done with *Chi square* test, comparing observed and expected frequencies, or with Student’s *t-test* (GraphPad QuickCalcs). To compare laser ablation measurements a parametric Student’s *t-test* was used. P-values of *Chi square* and *Student’s t* test are represented as: *< 0.05, **< 0.01, ***< 0.001 and ****< 0.0001.

## Acknowledgements

We thank the BDSC for reagents. This work was supported by the Spanish Agencia Estatal de Investigación (MCUI/AEI, http://www.ciencia.gob.es/; grant numbers PID2019-109013GB-I00 to MDM-B, PID2024-155234 NB-I00 to AG-R and CEX2020-001088-M to MDM-B and AG-R), by the Junta de Andalucía (grant number P20_00888 to MDM-B and AG-R) and by the European Regional Development Fund (http://ec.europa.eu/regional_policy/en/funding/erdf/). We thank the Consejería de Universidad, Investigación e Innovación, Junta de Andalucía, for financial support. Core funding to the CABD from the Junta de Andalucía is acknowledged.

## Author contributions

LR-O, CHF-E designed research, performed experiments, analyzed data and contributed to the writing of the manuscript. IMP performed experiments and provided useful comments on the manuscript. AG-R provided financial support, designed research, analyzed data and contributed to the writing of the manuscript. MDM-B provided financial support, performed experiments, designed research, analyzed data and wrote the manuscript.

## Competing interests

The authors declare no competing interests.

## Supplementary Information

**Supplementary Figure 1. Frequency and size of the multilayering phenotype increases with time.**

(A) Control and (B) mosaic egg chambers containing *mys^-^* FCs at the indicated stages, stained to visualize DNA (white) and GFP (green). Mutant FCs are identified by the absence of GFP signal. (C) Quantification of the multilayering phenotype at both anterior and posterior poles in mosaic egg chambers carrying *mys^-^* FCs. Scale bars, 20 μm.

**Supplementary Figure 2. Downregulation of laminin or collagen IV levels rescue the multilayering phenotype caused by clonal loss of integrin function.**

(A-F) S9 egg chambers of the indicated genotypes stained with the DNA marker Hoechst (white), anti-GFP (green) and Rhodamine Phalloidin (to label F-actin. RhPh, magenta). Mutant FCs are identified by the absence of GFP signal. (G) Quantification of the multilayering phenotype in egg chambers of the indicated genotypes. Scale bars, 20 μm.

## Supplemental Movies

**Movie S1. Spindle orientation during planar division of a control FC located at the posterior pole.**

Time-lapse movie of a control FC located at the posterior pole undergoing planar division. Centrosomes are labelled with *asl:*YFP (yellow), microtubules with *αtub:*GFP (green) and DNA with *his:*RFP (red). Scale bar, 3µm.

**Movie S2. Spindle orientation during non-planar division of a control FC located at the posterior pole.**

Time-lapse movie of a control FC located at the posterior pole undergoing non-planar division. Centrosomes are labelled with *asl:*YFP (yellow), microtubules with *αtub:*GFP (green) and DNA with *his:*RFP (red). Scale bar, 3µm.

**Movie S3. Non-planar division and reintegration of a control FC at the posterior pole.** Time-lapse movie of a control FC located at the posterior pole undergoing non-planar division. DNA is labelled with *his2AV:GFP* and *his:*RFP. Scale bar, 3µm.

**Movie S4. Non-planar division and reintegration of a control FC at the main body.**

Time-lapse movie of a control FC located at the main body undergoing non-planar division. DNA is labelled with *his2AV:GFP* and *his:*RFP. Scale bar, 3µm.

**Movie S5. Non-planar division and reintegration of a *mys* mutant FC at the main body.** Time-lapse movie of a *mys* mutant FC located at the main body undergoing non-planar division. DNA is labelled with *his:*RFP. Scale bar, 3µm.

**Movie S6. Non-planar division and reintegration of a *mys* mutant FC at the posterior pole of a monolayered epithelium.**

Time-lapse movie of a *mys* mutant FC located at the posterior pole of a monolayered epithelium undergoing non-planar division. DNA is labelled with *his:*RFP. Scale bar, 3µm.

**Movie S7. Non-planar division and reintegration of a *mys* mutant FC at the posterior pole of a multilayered epithelium.**

Time-lapse movie of a *mys* mutant FC located at the posterior pole of a multilayered epithelium undergoing non-planar division. DNA is labelled with *his:*RFP. Scale bar, 3µm.

**Movie S8. Laser ablation of cell bonds between control FCs**

Movie corresponds to the ablation experiment shown in Figure 4. The membranes of FCs are visualised with *resille:GFP*. A cell bond between two control FCs is ablated. Scale bar, 3µm. GFP fluorescent is lost in the middle of the ablated bond upon laser ablation. The movie continues 10s after the cut and shows displacement of the vertexes. Images are taken every 0.5 seconds.

**Movie S9. Planar division of a FC at the posterior pole expressing a *mys*RNAi.**

Time-lapse movie of a FC expressing a *mys RNAi*, located at the posterior pole, undergoing planar division. Cell membrane is labelled with *resille:GFP*. Scale bar, 3µm.

**Movie S10. Non-planar division of a FC at the posterior pole expressing a *mys*RNAi.** Time-lapse movie of a FC expressing a *mys RNAi*, located at the posterior pole, undergoing non-planar division. Cell membrane is labelled with *resille:GFP*. Scale bar, 3µm.

## References

1. Abdelilah-Seyfried, S., D.N. Cox, and Y.N. Jan. 2003. Bazooka is a permissive factor for the invasive behavior of *discs large* tumor cells in *Drosophila* ovarian follicular epithelia. Development. 130:1927–1935.

2. Bergstralh, D.T., H.E. Lovegrove, and D. St Johnston. 2013. Discs large links spindle orientation to apical-basal polarity in Drosophila epithelia. Curr Biol. 23:1707–1712.

3. Bergstralh, D.T., H.E. Lovegrove, and D. St Johnston. 2015. Lateral adhesion drives reintegration of misplaced cells into epithelial monolayers. Nat Cell Biol. 17:1497–1503.

4. Bilder, D. 2004. Epithelial polarity and proliferation control: links from the Drosophila neoplastic tumor suppressors. Genes Dev. 18:1909–1925.

5. Brakebusch, C., R. Grose, F. Quondamatteo, A. Ramirez, J.L. Jorcano, A. Pirro, M. Svensson, R. Herken, T. Sasaki, R. Timpl, S. Werner, and R. Fassler. 2000. Skin and hair follicle integrity is crucially dependent on beta 1 integrin expression on keratinocytes. EMBO J. 19:3990–4003.

6. Brodland, G.W. 2002. The Differential Interfacial Tension Hypothesis (DITH): a comprehensive theory for the self-rearrangement of embryonic cells and tissues. J Biomech Eng. 124:188–197.

7. Brown, N.H. 2000. Cell-cell adhesion via the ECM:integrin genetics in fly and worm. Matrix Biol. 19:191–201.

8. Bunch, T.A., and D.L. Brower. 1992. Drosophila PS2 integrin mediates RGD-dependent cell-matrix interactions. Development. 116:239–247.

9. Byrd, K.M., K.J. Lough, J.H. Patel, C.P. Descovich, T.A. Curtis, and S.E. Williams. 2016. LGN plays distinct roles in oral epithelial stratification, filiform papilla morphogenesis and hair follicle development. Development. 143:2803–2817.

10. Calvi, B.R., M.A. Lilly, and A.C. Spradling. 1998. Cell cycle control of chorion gene amplification. Genes Dev. 12:734–744.

11. Chen, J., and M.A. Krasnow. 2012. Integrin Beta 1 suppresses multilayering of a simple epithelium. PLoS One. 7:e52886.

12. Chou, T.B., and N. Perrimon. 1992. Use of a yeast site-specific recombinase to produce female germline chimeras in *Drosophila*. Genetics. 131:643–653.

13. Ciruna, B., A. Jenny, D. Lee, M. Mlodzik, and A.F. Schier. 2006. Planar cell polarity signalling couples cell division and morphogenesis during neurulation. Nature. 439:220–224.

14. Combedazou, A., V. Choesmel-Cadamuro, G. Gay, J. Liu, L. Dupre, D. Ramel, and X. Wang. 2017. Myosin II governs collective cell migration behaviour downstream of guidance receptor signalling. J Cell Sci. 130:97–103.

15. Crest, J., A. Diz-Munoz, D.Y. Chen, D.A. Fletcher, and D. Bilder. 2017. Organ sculpting by patterned extracellular matrix stiffness. Elife. 6.

16. Damen, M., L. Wirtz, E. Soroka, H. Khatif, C. Kukat, B.D. Simons, and H. Bazzi. 2021. High proliferation and delamination during skin epidermal stratification. Nature communications. 12:3227.

17. Diaz de la Loza, M.C., A. Diaz-Torres, F. Zurita, A.E. Rosales-Nieves, E. Moeendarbary, K. Franze, M.D. Martin-Bermudo, and A. Gonzalez-Reyes. 2017. Laminin Levels Regulate Tissue Migration and Anterior-Posterior Polarity during Egg Morphogenesis in Drosophila. Cell Rep. 20:211–223.

18. Dillon, M.E., L.R. Cahn, and R.B. Huey. 2007. Life history consequences of temperature transients in Drosophila melanogaster. J Exp Biol. 210:2897–2904.

19. Duffy, J.B., D.A. Harrison, and N. Perrimon. 1998. Identifying loci required for follicular patterning using directed mosaics. Developmen. 125:2263–2271.

20. Ewald, A.J., A. Brenot, M. Duong, B.S. Chan, and Z. Werb. 2008. Collective epithelial migration and cell rearrangements drive mammary branching morphogenesis. Dev Cell. 14:570–581.

21. Farhadifar, R., J.C. Roper, B. Aigouy, S. Eaton, and F. Julicher. 2007. The influence of cell mechanics, cell-cell interactions, and proliferation on epithelial packing. Curr Biol. 17:2095–2104.

22. Fernández-Miñán, A., M.D. Martín-Bermudo, and A. Gonzéalez-reyes. 2007. Integrin signaling regulates spindle orientation in *Drosophila* to preserve the follicular-epithelium monolayer. Current Biology. 17:683–688.

23. Fiore, V.F., M. Krajnc, F.G. Quiroz, J. Levorse, H.A. Pasolli, S.Y. Shvartsman, and E. Fuchs. 2020. Mechanics of a multilayer epithelium instruct tumour architecture and function. Nature. 585:433–439.

24. Goode, S., and N. Perrimon. 1997. Inhibition of patterned cell shape change and cel invasion by Discs large during *Drosophila* oogenesis. Genes and Development. 11:2532–2544.

25. Gutzeit, H.O., W. Eberhardt, and E. Gratwohl. 1991. Laminin and basement membrane-associated microfilaments in wild type and mutant *Drosophila* ovarian follicles. J. Cell Science. 100:781–788.

26. Jones, P.H., and F.M. Watt. 1993. Separation of human epidermal stem cells from transit amplifying cells on the basis of differences in integrin function and expression. Cell. 73:713–724.

27. King, R.C. 1970. Ovarian development in Drosophila melanogaster. Academic Press, New York, NY.

28. Lechler, T., and E. Fuchs. 2005. Asymmetric cell divisions promote stratification and differentiation of mammalian skin. Nature. 437:275–280.

29. Lee, J.K., E. Brandin, D. Branton, and L.S. Goldstein. 1997. alpha-Spectrin is required for ovarian follicle monolayer integrity in Drosophila melanogaster. Development. 124:353–362.

30. Lee, S.B., K.S. Cho, E. Kim, and J. Chung. 2003. blistery encodes Drosophila tensin protein and interacts with integrin and the JNK signaling pathway during wing development. Development. 130:4001–4010.

31. Li, M.A., J.D. Alls, R.M. Avancini, K. Koo, and D. Godt. 2003. The large Maf factor Traffic Jam controls gonad morphogenesis in Drosophila. Nat Cell Biol. 5:994–1000.

32. Lovegrove, H.E., D.T. Bergstralh, and D. St Johnston. 2019. The role of integrins in Drosophila egg chamber morphogenesis. Development. 146.

33. Martin-Bermudo, M.D., and N.H. Brown. 1999. Uncoupling integrin adhesion and signaling: the b_PS_ cytoplasmic domain is sufficient to regulate gene expression in the *Drosophila* embryo. Genes Dev. 13:729–739.

34. Meignin, C., I. Alvarez-Garcia, I. Davis, and I.M. Palacios. 2007. The salvador-warts-hippo pathway is required for epithelial proliferation and axis specification in Drosophila. Curr Biol. 17:1871–1878.

35. Miroshnikova, Y.A., H.Q. Le, D. Schneider, T. Thalheim, M. Rubsam, N. Bremicker, J. Polleux, N. Kamprad, M. Tarantola, I. Wang, M. Balland, C.M. Niessen, J. Galle, and S.A. Wickstrom. 2018. Adhesion forces and cortical tension couple cell proliferation and differentiation to drive epidermal stratification. Nat Cell Biol. 20:69–80.

36. Molina Lopez, E., A. Kabanova, A. Winkel, K. Franze, I.M. Palacios, and M.D. Martin-Bermudo. 2023. Constriction imposed by basement membrane regulates developmental cell migration. PLoS Biol. 21:e3002172.

37. Morin, X., R. Daneman, M. Zavortink, and W. Chia. 2001. A protein trap strategy to detect GFP-tagged proteins expressed from their endogenous loci in Drosophila. Proc Natl Acad Sci U S A. 98:15050–15055.

38. Moyle, L.A., R.Y. Cheng, H. Liu, S. Davoudi, S.A. Ferreira, A.A. Nissar, Y. Sun, E. Gentleman, C.A. Simmons, and P.M. Gilbert. 2020. Three-dimensional niche stiffness synergizes with Wnt7a to modulate the extent of satellite cell symmetric self-renewal divisions. Mol Biol Cell. 31:1703–1713.

39. Ng, B.F., G.K. Selvaraj, C. Santa-Cruz Mateos, I. Grosheva, I. Alvarez-Garcia, M.D. Martin-Bermudo, and I.M. Palacios. 2016. alpha-Spectrin and integrins act together to regulate actomyosin and columnarization, and to maintain a monolayered follicular epithelium. Development. 143:1388–1399.

40. Packard, A., K. Georgas, O. Michos, P. Riccio, C. Cebrian, A.N. Combes, A. Ju, A. Ferrer-Vaquer, A.K. Hadjantonakis, H. Zong, M.H. Little, and F. Costantini. 2013. Luminal mitosis drives epithelial cell dispersal within the branching ureteric bud. Dev Cell. 27:319–330.

41. Petridou, N.I., and P.A. Skourides. 2016. A ligand-independent integrin beta1 mechanosensory complex guides spindle orientation. Nature communications. 7:10899.

42. Polesello, C., and N. Tapon. 2007. Salvador-warts-hippo signaling promotes Drosophila posterior follicle cell maturation downstream of notch. Curr Biol. 17:1864–1870.

43. Santa-Cruz Mateos, C., A. Valencia-Exposito, I.M. Palacios, and M.D. Martin-Bermudo. 2020. Integrins regulate epithelial cell shape by controlling the architecture and mechanical properties of basal actomyosin networks. PLoS Genet. 16:e1008717.

44. Schaeffer, V., C. Althauser, H.R. Shcherbata, W.M. Deng, and H. Ruohola-Baker. 2004. Notch-dependent Fizzy-related/Hec1/Cdh1 expression is required for the mitotic-to-endocycle transition in Drosophila follicle cells. Curr Biol. 14:630–636.

45. Seldin, L., A. Muroyama, and T. Lechler. 2016. NuMA-microtubule interactions are critical for spindle orientation and the morphogenesis of diverse epidermal structures. Elife. 5.

46. Simpson, C.L., D.M. Patel, and K.J. Green. 2011. Deconstructing the skin: cytoarchitectural determinants of epidermal morphogenesis. Nat Rev Mol Cell Biol. 12:565–580.

47. Spradling, A.C. 1993. Developmental genetics of oogenesis. The Development of Drosophila melanogaster. M. Bate and A Martinez-Arias, editors. Cold Spring Harbor Lab. Press, Cold Spring Harbor, New York:1–70.

48. Tanentzapf, G., C. Smith, J. McGlade, and U. Tepass. 2000. Apical, Lateral, and Basal polarization cues contribute to the development of the follicular epithelium during *drosophila* oogenesis. J. Cell Biology. 151:891–904.

49. Valencia-Exposito, A., M.J. Gomez-Lamarca, T.J. Widmann, and M.D. Martin-Bermudo. 2022. Integrins Cooperate With the EGFR/Ras Pathway to Preserve Epithelia Survival and Architecture in Development and Oncogenesis. Front Cell Dev Biol. 10:892691.

50. Yee, G.H., and R.O. Hynes. 1993. A novel, tissue-specific integrin subunit, b_g_, expressed in the midgut of *Drosophila melanogaster*. Development. 118:845–858.

